# Dynamic changes in the transcriptome of oocytes during adolescent-onset PCOS in mice

**DOI:** 10.1101/2022.11.08.515666

**Authors:** Du Danfeng, Deng Ke, Fan Dengxuan, Xu Congjian

**Author notes:** Xu Congjian is the **Post-Publication Corresponding author** and the **Last author**. Du Danfeng is the **First author** and will **handle correspondence** at all stages of refereeing and publication, Deng Ke contributed equally to this work and should be listed as **Co-first author**. First author. Co-First author. Second author. Zip code: 200000.

## Abstract

(1) Background: This study aimed to explore temporal changes in the transcriptome of oocytes in an adolescent-onset polycystic ovarian syndrome (PCOS) mouse model. (2) Methods: An adolescent-onset PCOS mouse model was established using DHEA. Genes with a similar expression trend over time were identified using trend analysis. KEGG pathway enrichment analysis and gene regulatory network diagrams were examined for signaling pathways to identify potential hub genes related to the pathogenesis of PCOS. (3) Results: Four main trends of gene expression were extracted, of which six combinations of Venn diagrams were generated. Differentially expressed genes were mainly enriched in oxidative phosphorylation, cell cycle, P53 signaling pathway. Cell cycle-related genes (Skp1, Ccnb1, Orc1 and 5, Wee2, Mapk3, Cdc20) were abnormally down-regulated in the DHEA group. Ptges3 was the top1 DEGs at the initial stage of PCOS modeling. (4) Conclusion: This study provides a novel insight into the altered transcriptome of oocytes from PCOS mice. mtDNA-related genes and Cell cycle-related genes play the most important role in the development of PCOS. Ptges3 was the one of the top DEGs which was up-regulated in DHEA group at the initial stage of modeling, which suggested it may play an important role in the early stage of PCOS.

## 1. Introduction

Polycystic ovarian syndrome (PCOS) is a common endocrine and metabolic dysfunction. Patients with PCOS may have an increased number of retrieved oocytes during IVF treatment, but the embryo cleavage rate is low for these patients [1]. Embryo developmental competence was reduced in a PCOS model induced by dehydroepiandrosterone (DHEA) [2]. The pathogenesis of PCOS is not fully elucidated, but some evidence indicates a strong heritable component in the development of PCOS. Several genome-wide association studies have identified approximately 200 candidate genes related to the risk of PCOS [3], but not a single fully penetrant variant gene has been reported to date. All candidate genes or mutations show a low penetrance associated with many other covariant factors, such as hormones, environmental changes or lifestyles [4].

Considerable work has been done to classify PCOS in the last several decades, but the consensus regarding PCOS phenotypes remains controversial. Some researchers propose PCOS is mainly caused by ovarian-derived androgen excess, while others suggest it is a complex systemic metabolism disorder [5]. Fetal exposure to excessive androgen, nutritional excess and hyperglycemia during critical stages of prenatal development may program neurobiological circuitry [6]. Prenatally androgenized androgen receptor knockout mice and neuron-specific androgen receptor knockout mice did not develop PCOS-like symptoms [7].

PCOS can be divided into adolescent- and adult-onset types according to the time of onset. During the pubertal maturation process of the hypothalamic-pituitary-ovarian axis, girls with a predisposition to PCOS may exhibit increased gonadotropin-releasing hormone and luteinizing hormone pulse frequencies, ultimately leading to an increased luteinizing hormone/follicle stimulating hormone ratio [8]. Some characteristics of PCOS, such as acne, irregular menses and polycystic ovary morphology, may also be normal physiological symptoms of puberty [9]. Puberty may activate the susceptibility to develop PCOS, which interferes with the normal pubertal transition. Other factors, such as elevated anti-Mullerian hormone secretion, luteinizing hormone secretion and gonadotropin-releasing hormone pulse frequency, may disrupt the pubertal transition [10]. Girls who do not establish a normal hypothalamic-pituitary-ovarian axis and mature along the PCOS trajectory may eventually develop persistent PCOS [11].

At present, the origin theory of PCOS is more inclined to the ‘adolescence origin or developmental origin’, in which adult-onset disease originates from maternal exposure and a high androgen environment or the abnormal establishment of hypothalamic-pituitary-ovarian axis feedback mechanisms in puberty. Refractory or medicine-resistant PCOS usually arises during puberty and is marked by insulin resistance, hyperinsulinemia and hyperandrogenism [12]. Many polygenic genetic diseases may originate from maternal-derived abnormal genes in oocytes. Mouse models have long been used to study PCOS, yet despite many reports describing the reproductive and metabolic phenotype of PCOS mouse models, research on the oocyte itself remains sparse [13]. The incidence rate of PCOS is low in women older than 35 years, and adult-onset PCOS is more likely to be self-limited with aging or relieved by medical treatment. Most of the cases with drug resistant or familial-like PCOS develop their symptoms from puberty, and may be described as having adolescent-onset PCOS. The latter may be the type of PCOS that should be given more attention in the research and clinical setting. The aim of this study was to investigate the dynamic temporal changes of the oocyte transcriptome during PCOS modeling. Transcriptome data at six consecutive time points during PCOS mouse modeling were analyzed to detect the dynamic trend of gene regulation and reveal critical genes or pathways that may be important candidate factors for PCOS development and further studies.

## 2. Materials and Methods

### 2.1 Construction of the adolescent onset PCOS mouse model

The prepubertal mouse was used to mimic the hyperandrogenic phenotype of patients with adolescent onset PCOS. Female ICR mice were purchased from Shanghai JieSiJie Laboratory Animal Co. [production license number: SCXK (Shanghai) 2018-0004, China] and housed at the laboratory animal center of Obstetrics and Gynecology Hospital of Fudan University. Mice were housed with free access to food and water, and a constant temperature, humidity and light duration. At 25 days of age, mice were subcutaneously injected with DHEA powder (MCE, HY-14650, China) (6 mg/100 g body weight/d) dissolved in corn oil (MCE, HY-1888, China) (Figure1). All procedures were performed in accordance with the guidelines of Experimental Animals Management Regulations (version 2017-03-01, China).

### 2.2 Ovary morphology and serum testosterone measurement

Mice were executed by carbon dioxide suffocation method. Ovaries of mice from each group were removed quickly and fixed in 4% paraformaldehyde for morphological assessment. Five-μm sections of ovary tissue were mounted on glass slides and stained with hematoxylin and eosin. Ovary morphology was further examined under light microscopy. The day after the final DHEA injection, blood was collected and centrifugated at a speed of 1000×g for 10 min. Serum testosterone was measured using a mouse testosterone enzyme immunoassay kit (Westang Bio-tech Co., Ltd., F11580, China).

### 2.3 Oocyte collection

From 25 days of age to two weeks after the final DHEA injection, oocytes were collected every other week so that six time points were established (points 0–5, Figure 1). At each time point, a mouse from each group was selected randomly, one ovary was fixed in paraformaldehyde for morphology assessment, and the other ovary was punctured with a 1 ml syringe needle in HEPES (Sigma, 83264, USA) to collect germinal vesicle-stage oocytes. Granulosa cells were removed mechanically. Ten oocytes of the same size, translucent and in good condition, were selected from each group for RNA sequencing.

**Figure1.**
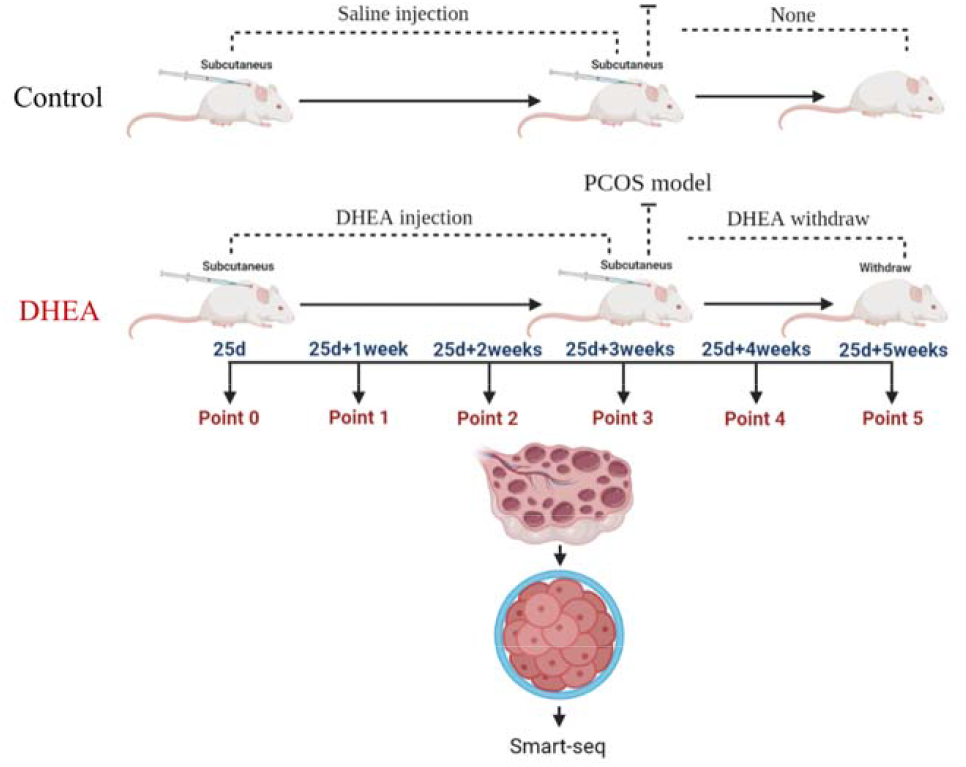
The schematics of the study design.

### 2.4 Transcriptome sequencing

The selected oocytes were processed for RNA extraction and reverse transcription within 1 h and were subjected to cDNA amplification and purification. Reverse transcription was performed using a SMARTer Ultra Low RNA Kit (Clontech, USA) directly in the cell lysate. Then, cDNA was amplified and purified using an Advantage 2 PCR Kit (Clontech). The cDNA library was sequenced using an Illumina sequencing platform (NovaSeq6000). The raw reads were cleaned by removing adaptor sequences, short sequences (length < 35 bp), low-quality bases (quality < 20) and ambiguous sequences. Hisat2 (version:2.0.4) was used to map the cleaned RNA-seq reads to the mouse mm10 genome with two mismatches, two gaps and one multi-hit allowed. The gene expression value was normalized by fragments per kilobase of exon per million mapped fragments (FPKM) and adjusted by a geometric algorithm (All the transcriptome sequencing method was provided by Shanghai Biotechnology Corporation, China, Contract number E20210017-5).

### 2.5 Immunofluorescence staining of oocytes

Oocytes were denuded mechanically and fixed in 1% paraformaldehyde for 30min. Then, oocytes were transferred into 0.2% Triton X-100 in phosphate buffered saline (PBS) to permeabilize for 30 min at room temperature. After blocking in QuickBlock blocking buffer (Beyotime, P0260, China) for 15 min, oocytes were incubated with Anti-Ptges3 antibody (1:250, ab92503, UK) overnight at 4°C, followed by three washes (5 min each) with PBS the next day. Oocytes were then incubated with secondary antibody (1:500 dilution of Alexa Fluor 488 goat anti-rabbit antibody, abcam, ab150077) for 1 h at room temperature protected from light. Oocytes were then mounted in a drop of antifade mounting medium (Beyotime, P0126) on a siliconized slide. Images were taken using a laser confocal microscope (Leica, Wetzlar, Germany).

### 2.6 Statistical Analysis

Data analysis was performed using the OmicShare tools, an open online platform for bioinformatics analysis (https://www.omicshare.com/tools, China) (Figure 2). At each time point (Point 1–5), two group of genes were compared to identify differentially expressed genes (DEGs). To further investigate dynamic gene expression patterns during the modeling process, trend analysis was used to determine genes with the same expression trend over six consecutive time points in the control and DHEA groups. Genes with very small expression levels were considered to play a limited role in the development of the disease, so the raw data were filtered by removing genes with FPKM < 1. Data of four main trends were extracted from each group and used for further analysis. Genes with the same or different expression trends from each group were compared for overlaps and differences, and the results were visualized using Venn diagrams. Genes with different or contrary expression trends from the Venn diagrams were considered to be abnormal and were used for gene pathway enrichment analysis (Kyoto Encyclopedia of Genes and Genomes, KEGG) (https://www.kegg.jp/)[14-16] and gene regulatory network analysis to explore the function of these genes and pathways. Genes from the critical pathway of each gene regulatory network were listed as hub genes. Data of relative grey value was analyzed in SPSS 19.0 (IBM Corp, USA). A value of *P* < 0.05 was considered statistically significant.

**Figure2.**
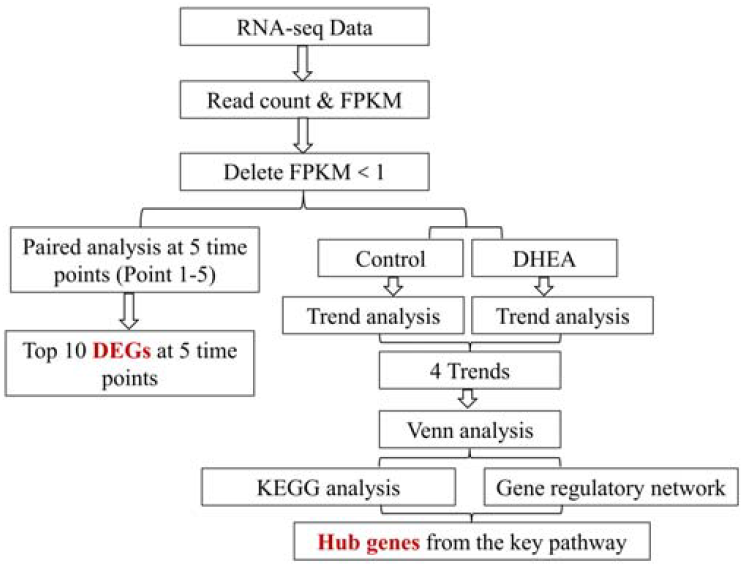
RNA-seq data analysis flowchart.

## 3. Results

### 3.1 Reproductive characteristics of PCOS mice

After three weeks of DHEA injection, the PCOS mouse model was successfully established (Figure 3A). Ovarian tissue histology showed large numbers of follicles at different stages of development in the control group, while ovaries in the PCOS group exhibited significantly increased cystic follicles with degenerate granulosa cell layers. Serum testosterone levels were significantly higher in the PCOS group versus healthy controls (Figure 3B). After the final DHEA injection, ovarian morphology gradually recovered to a normal state within 3 weeks (Figure 3C).

**Figure3.**
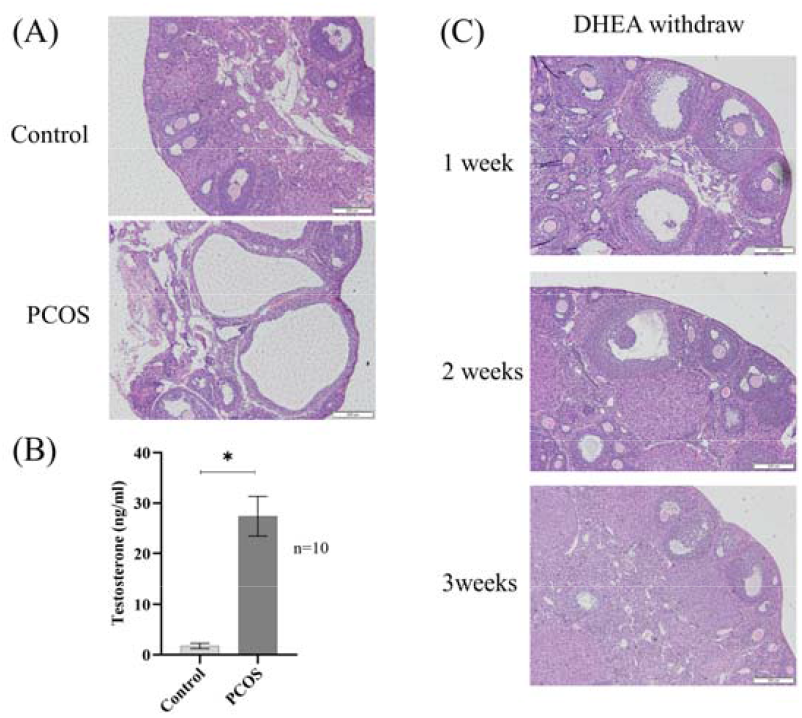
Ovarian histology and serum testosterone. (a) HE staining of ovarian sections from both control and PCOS model (point3); (b) Serum testosterone level of both groups at time point3; (C) HE staining of ovarian sections from DHEA group after DHEA injection stopped for 1, 2 and 3 weeks.

### 3.2 DEGs at different time points

Data from two groups were compared to identify genes up- or down-regulated in the DHEA group at each of the five time points examined (points 1–5). The top ten DEGs are listed below and ranked by Log2FC (Table 1-4).

**Table 1.**
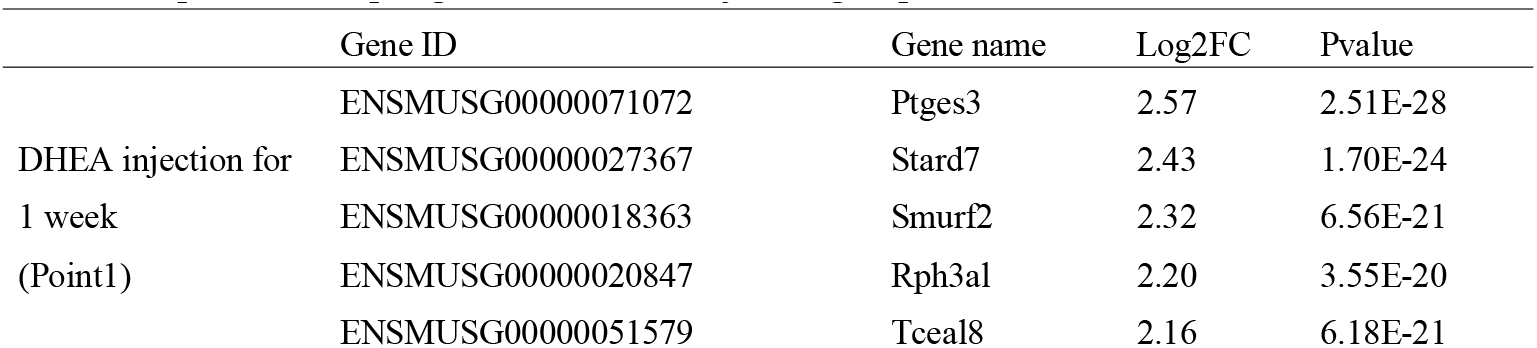
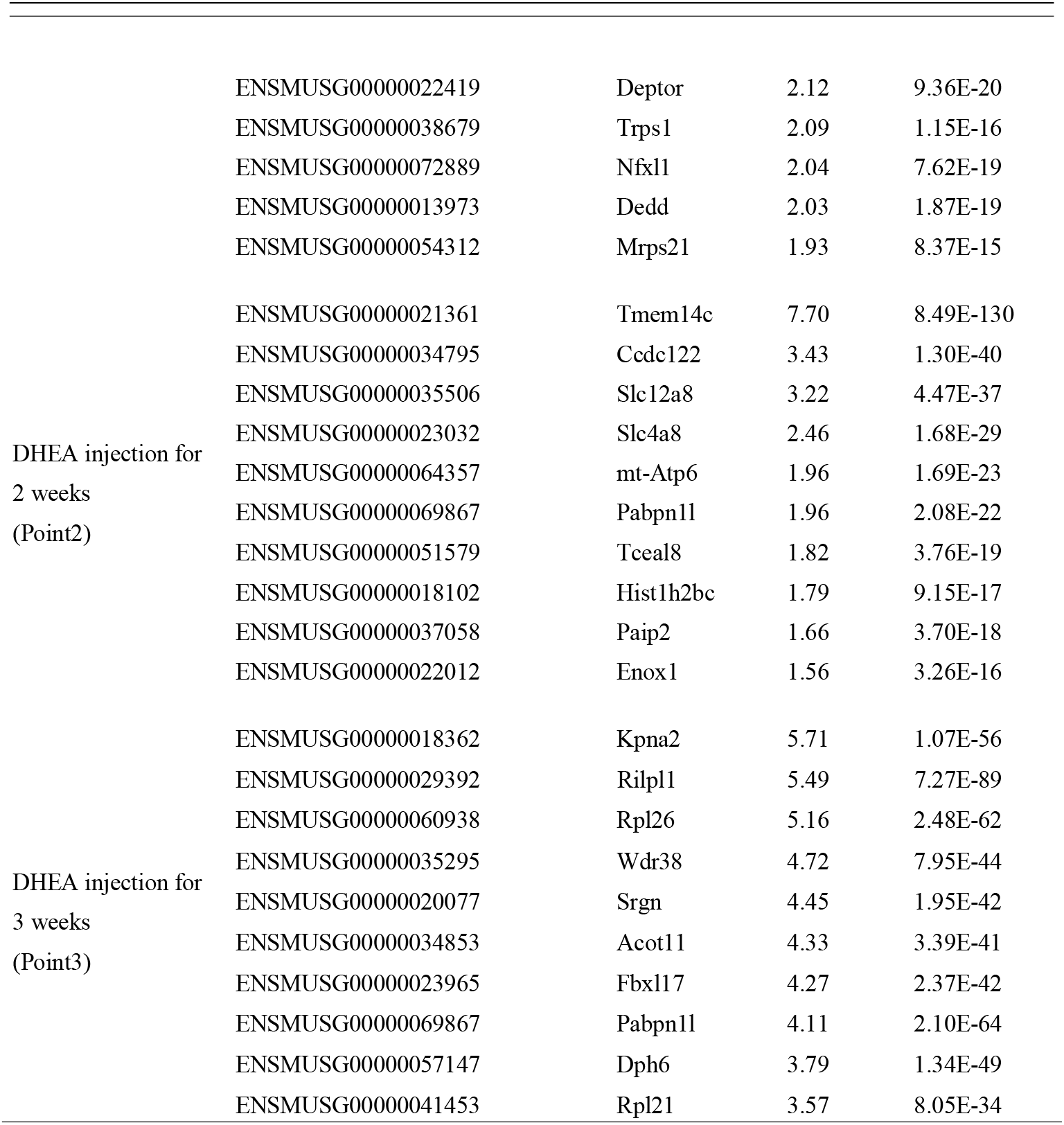
Top 10 DEGs Up-regulated in DHEA injection group.

**Table 2.**
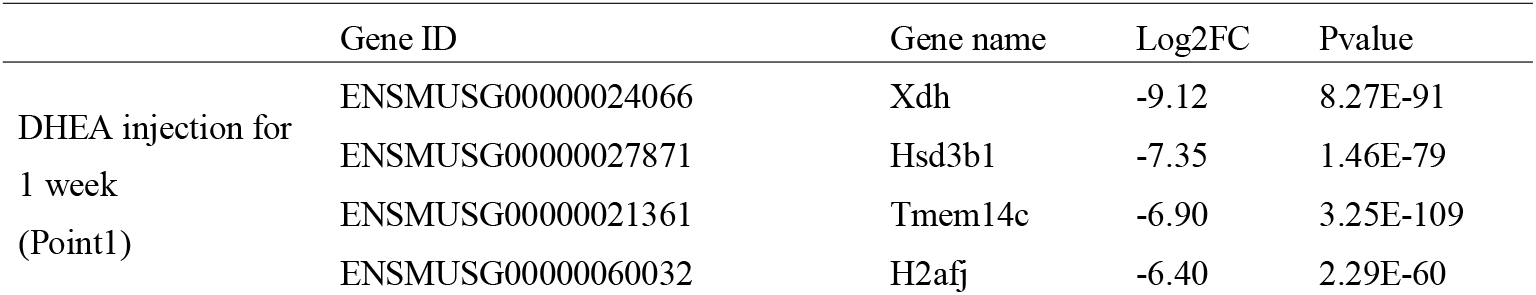
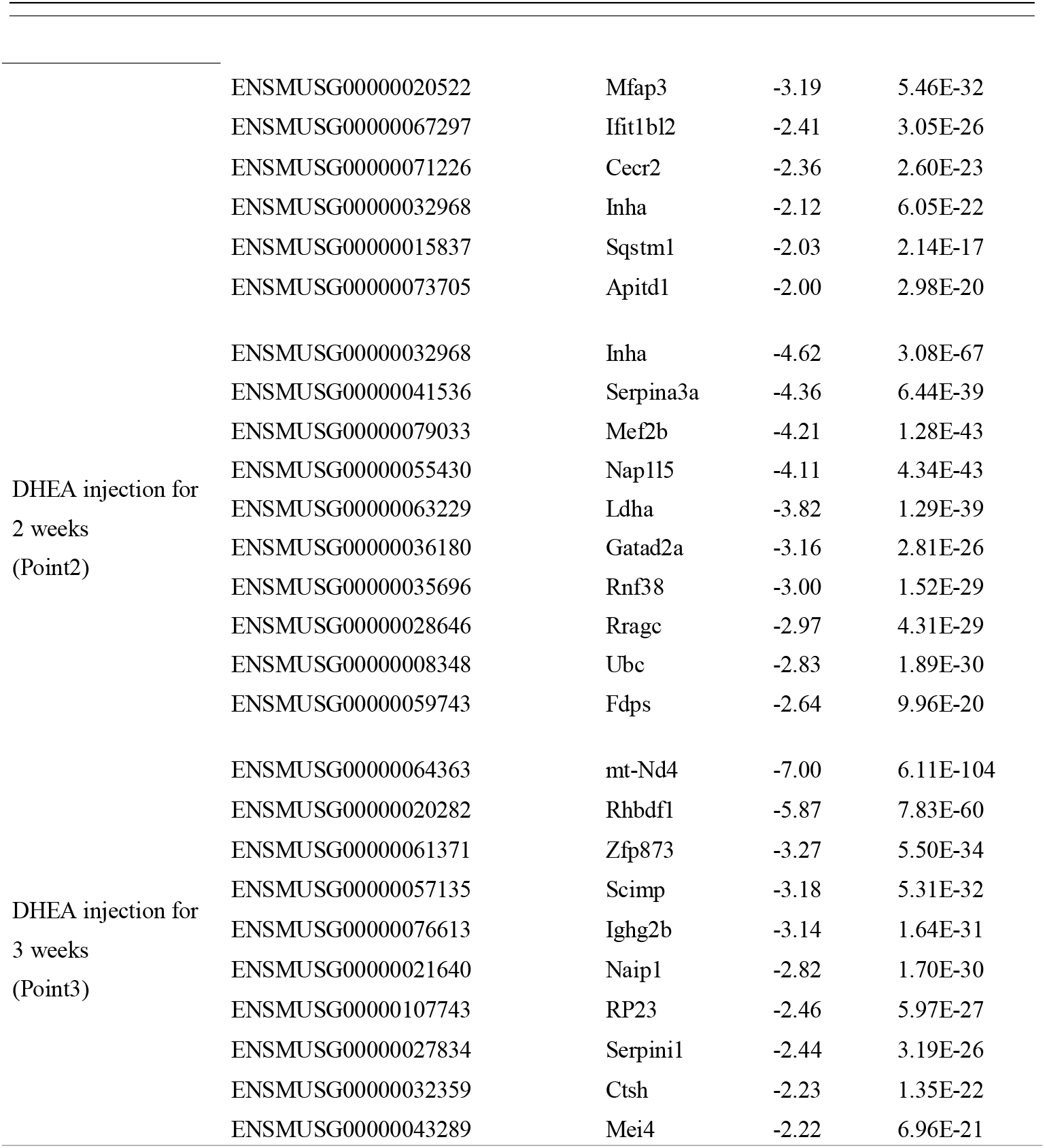
Top 10 DEGs Down-regulated in DHEA injection group.

**Table 3.**
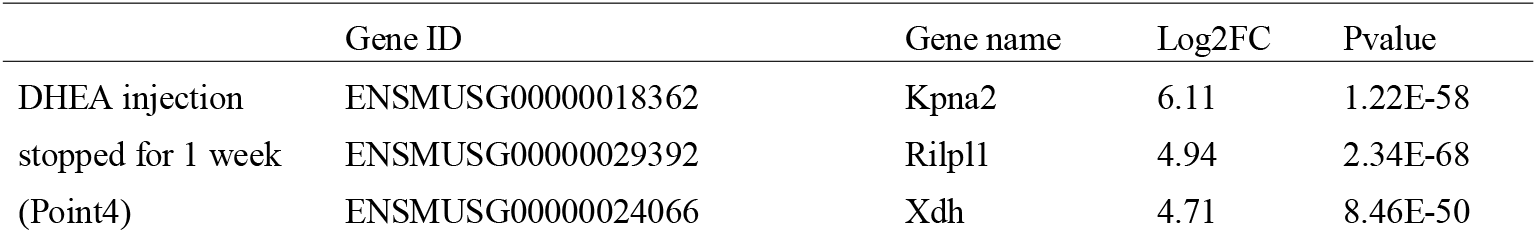
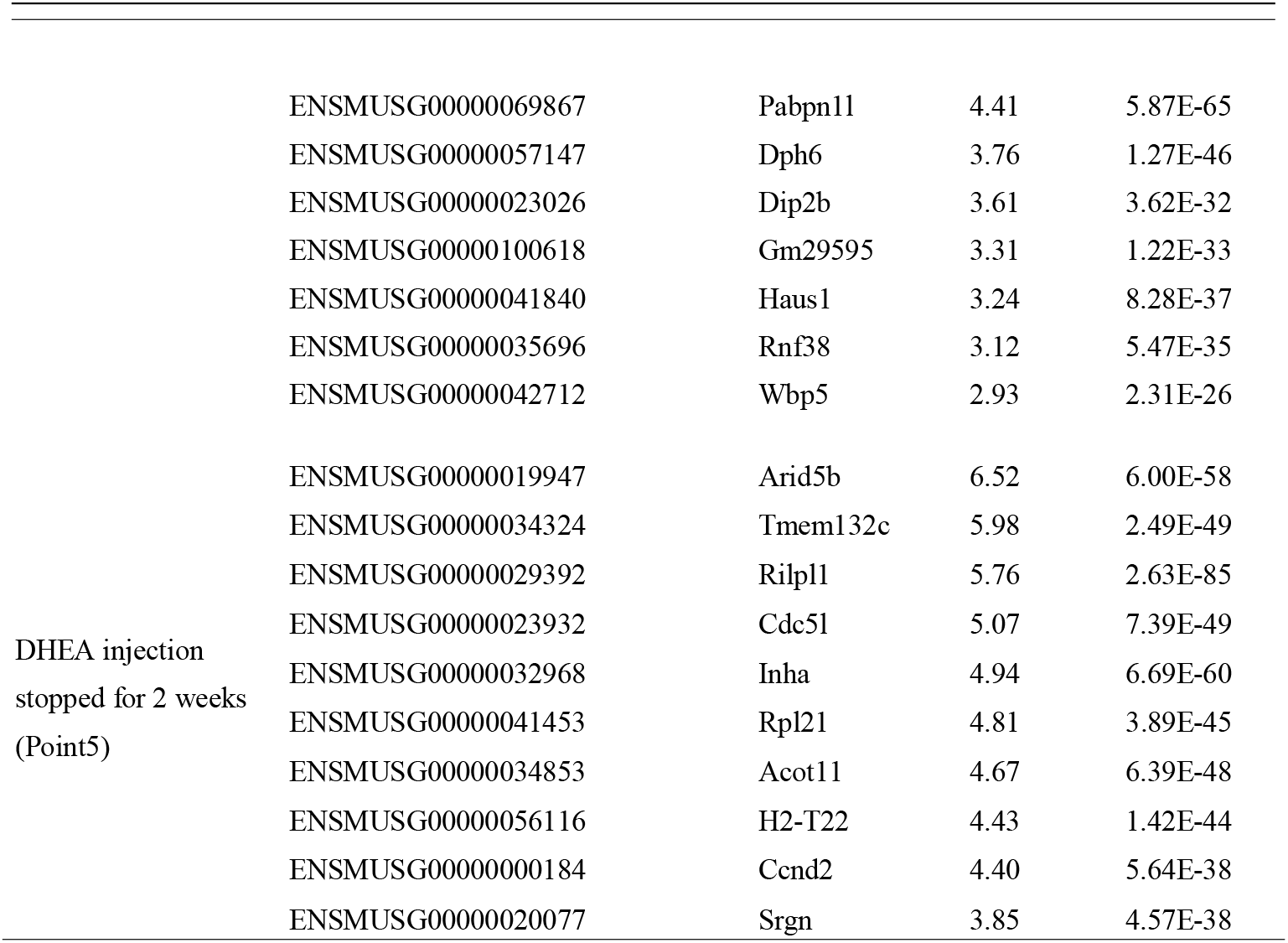
Top 10 DEGs Up-regulated in DHEA group.

**Table 4.**
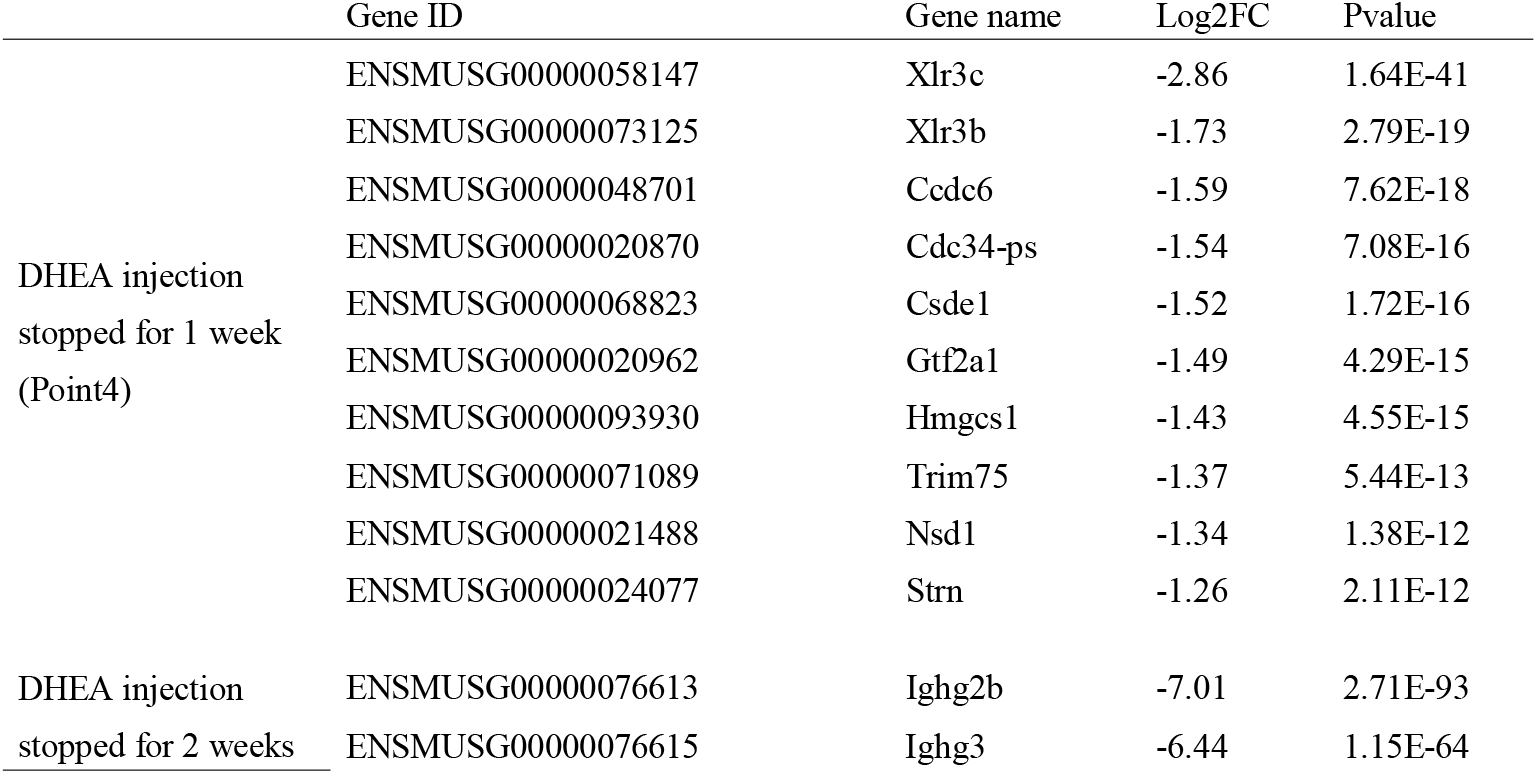
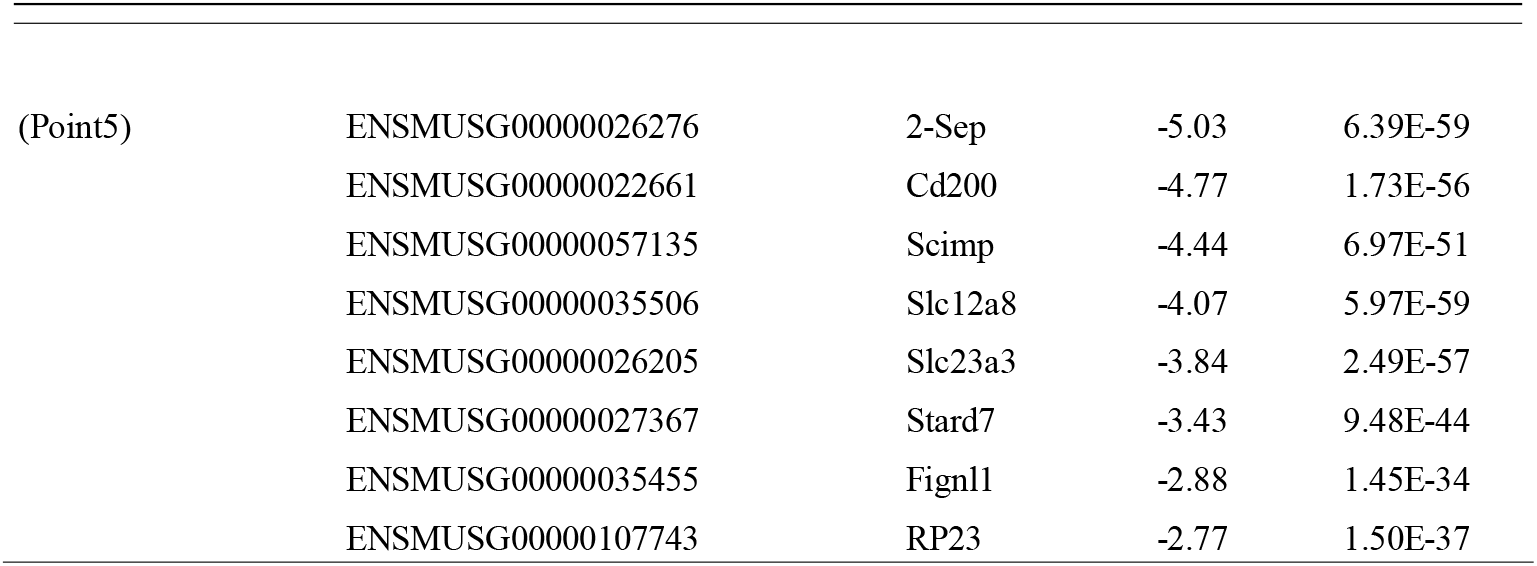
Top 10 DEGs Down-regulated in DHEA group.

### 3.3 Trend analysis

A total of 21,928 genes were identified in mouse oocytes, and 9,382 genes had FPKM > 1. Given that each gene had a different trend over time, data in each group was first analyzed through trend analysis to summarize genes with similar trends over time. Four main trends of data were selected for further analysis. In the control group, analysis over six consecutive time points (points 0–5) showed 314 genes were continuously up-regulated, 525 genes were continuously down-regulated, 717 genes had a down-flat-up regulation trend and 859 genes had an up-flat-down regulation trend (Figure 4A). In the DHEA group over the same time points, 1491 genes were continuously up-regulated, 811 genes were continuously down-regulated, 1222 genes showed a down-flat-up regulation trend and 334 genes showed an up-flat-down regulation trend (Figure 4B).

**Figure4.**
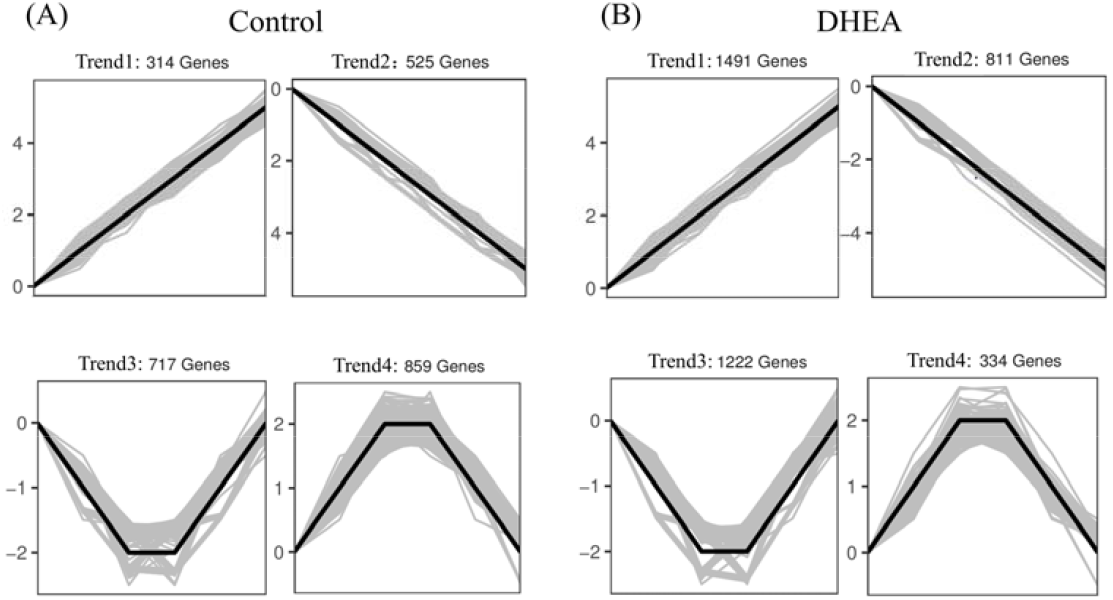
Trend analysis of each group. (a) In the control group, 314 genes were continuously up-regulated, 525 genes were continuously down-regulated, 717 genes had a down-flat-up r egulation trend and 859 genes had an up-flat-down regulation trend. (**b**) In the DHEA group, 1491 genes were continuously up-regulated, 811 genes were continuously down-regulated, 1222 genes showed a down-flat-up regulation trend and 334 genes showed an up-flat-down regulation trend.

### 3.4 Venn diagram analysis

Four selected trends of data were plotted as Venn diagrams to identify genes with abnormal expression trends in the DHEA group. For genes with the same trend in both groups, genes only expressed in the DHEA group were selected. For genes with a contrary trend, overlapping genes of the two groups were selected. Six kinds of data combinations and Venn diagrams were finally produced. Venn1 was produced with genes expressed as ‘trend1’ in both the control and DHEA groups (Figure 5A). In this diagram, 1372 genes were abnormally continuously up-regulated in the DHEA group (Figure 5B). Venn2 was produced with genes expressed as ‘trend2’ in both the control and DHEA groups (Figure 6A), with 743 genes abnormally down-regulated in the DHEA group(Figure 6B). Venn3 was produced with genes expressed as ‘trend1’ in the control group and genes expressed as ‘trend2’ in the DHEA group (Figure 7A). Twenty genes were abnormally down-regulated in the DHEA group (Figure 7B). Venn4 was produced with genes expressed as ‘trend2’ in the control group and genes expressed as ‘trend1’ in the DHEA group (Figure 8A). Seventy-one genes were abnormally up-regulated in the DHEA group (Figure 8B). Venn5 was produced with genes expressed as ‘trend3’ in both groups (Figure 9A), in which 967 genes were abnormally down-flat-up-regulated in the DHEA group (Figure 9B). Venn6 was produced with genes expressed as ‘trend4’ in both groups (Figure 10A), in which 190 genes were abnormally up-flat-down-regulated in the PCOS group (Figure 10B). The selected genes are marked with the red dotted circle.

**Figure5.**
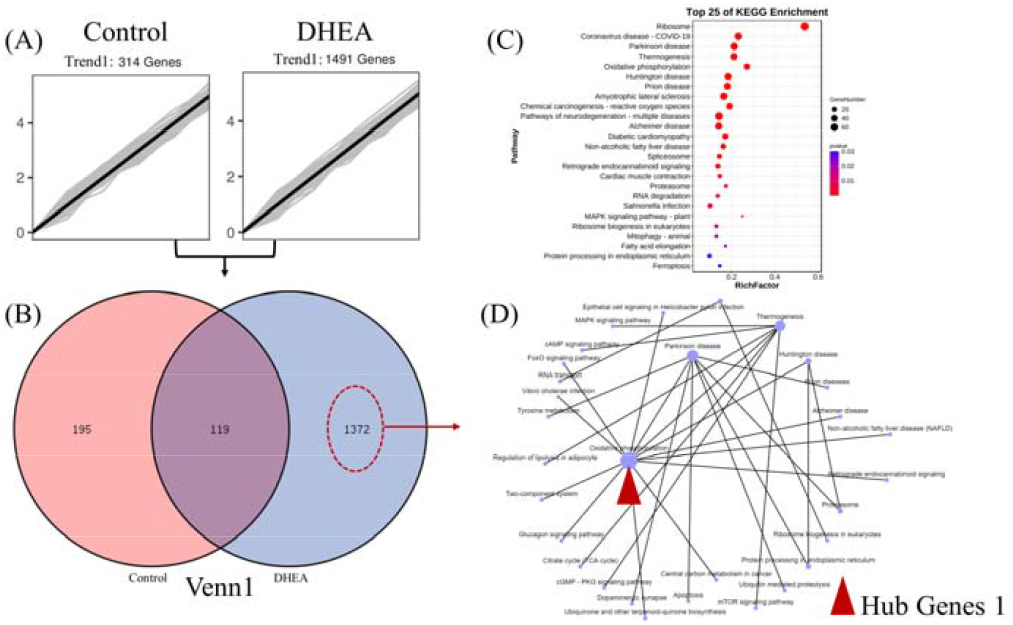
Analysis of genes expressed as ‘trend1’ in both groups. (a) 314 and 1491 genes were continuously up-regulated in each group separately. (b) Venn1 diagram showed 1372 genes were abnormally up-regulated in DHEA group. (c) KEGG analysis of the 1372 genes abnormally up-regulated in DHEA group. (d) Gene regulatory network diagram of the 1372 genes abnormally up-regulated in DHEA group.

**Figure6.**
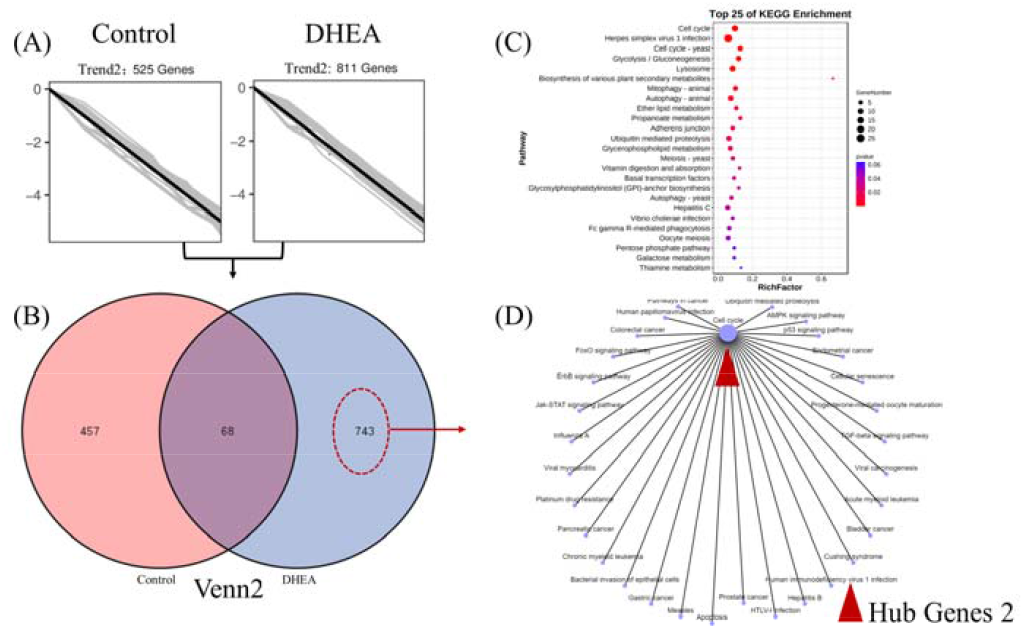
Analysis of genes expressed as ‘trend2’ in both groups. (a) 525 and 811 genes were continuously down-regulated in each group separately. (b) Venn2 diagram showed 743 genes were abnormally up-regulated in DHEA group. (c) KEGG analysis of the 743 genes abnormally down-regulated in DHEA group. (d) Gene regulatory network diagram of the 743 genes abnormally down-regulated in DHEA group.

**Figure7.**
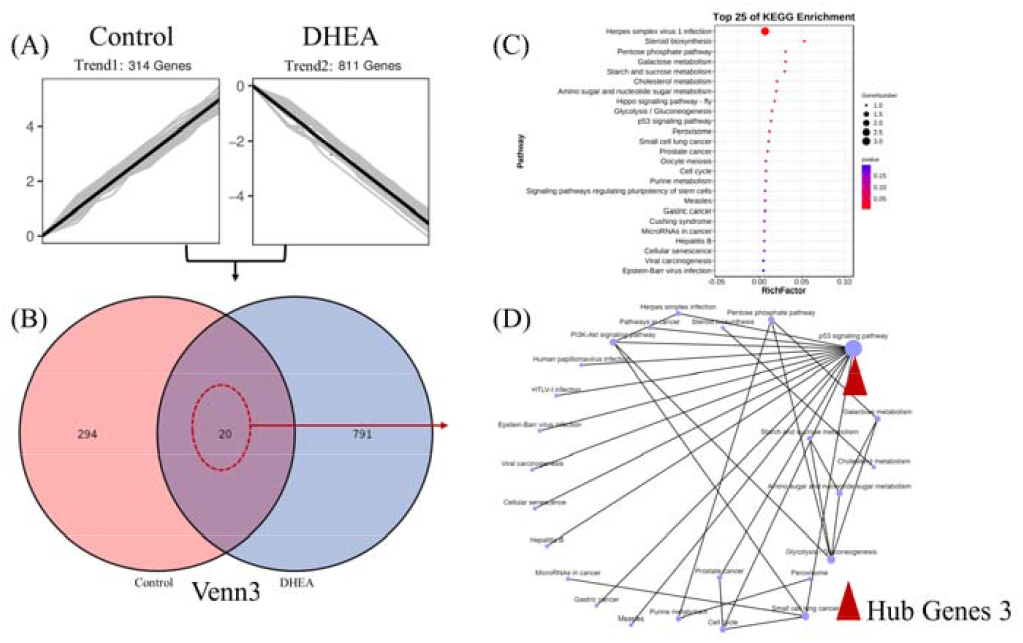
Analysis of genes expressed as ‘trend1’ in the control group and genes expressed as ‘trend2’ in the DHEA group. (a) 314 genes were up-regulated in control group and 811 genes were down-regulated in DHEA group. (b) Venn3 diagram showed 20 genes were abnormally down-regulated in DHEA group. (c) KEGG analysis of the 20 genes abnormally down-regulated in DHEA group. (d) Gene regulatory network diagram of the 20 genes abnormally down-regulated in DHEA group.

**Figure8.**
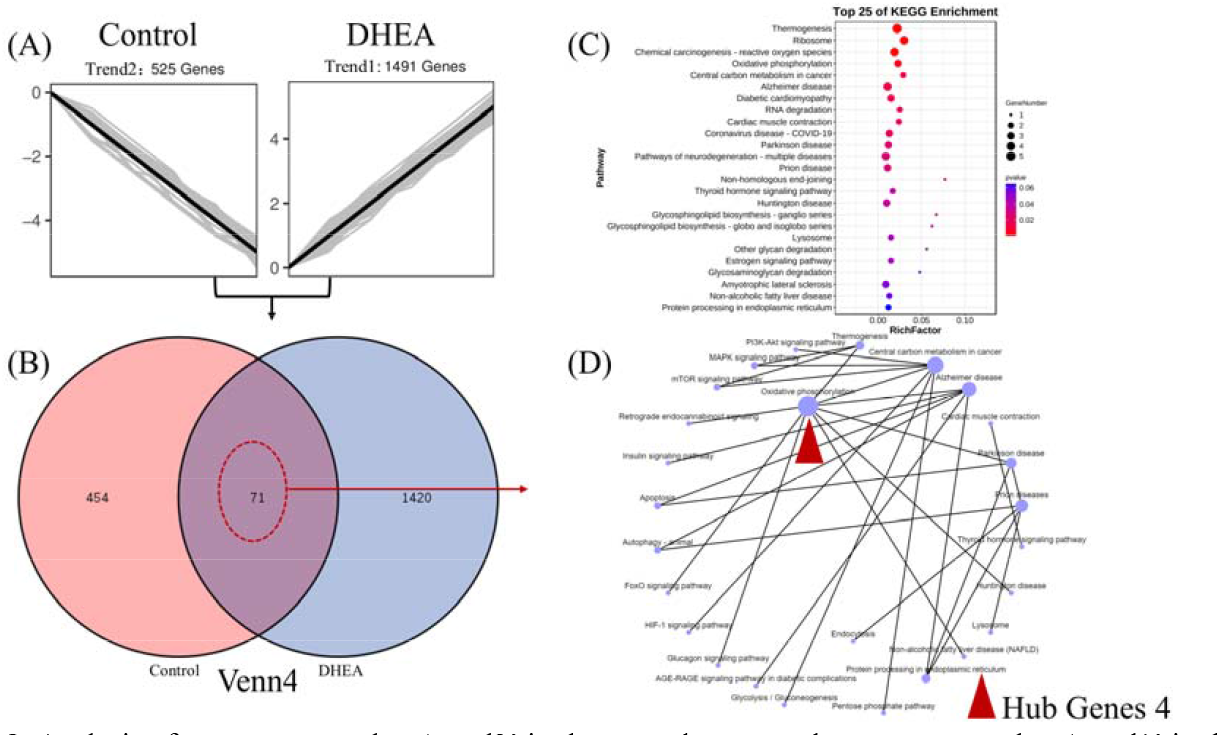
Analysis of genes expressed as ‘trend2’ in the control group and genes expressed as ‘trend1’ in the DHEA group. (a) 525 genes were down-regulated in control group and 1491 genes were up-regulated in DHEA group. (b) Venn4 diagram showed 71 genes were abnormally up-regulated in DHEA group. (c) KEGG analysis of the 71 genes abnormally up-regulated in DHEA group. (d) Gene regulatory network diagram of the 71 genes abnormally up-regulated in DHEA group

**Figure9.**
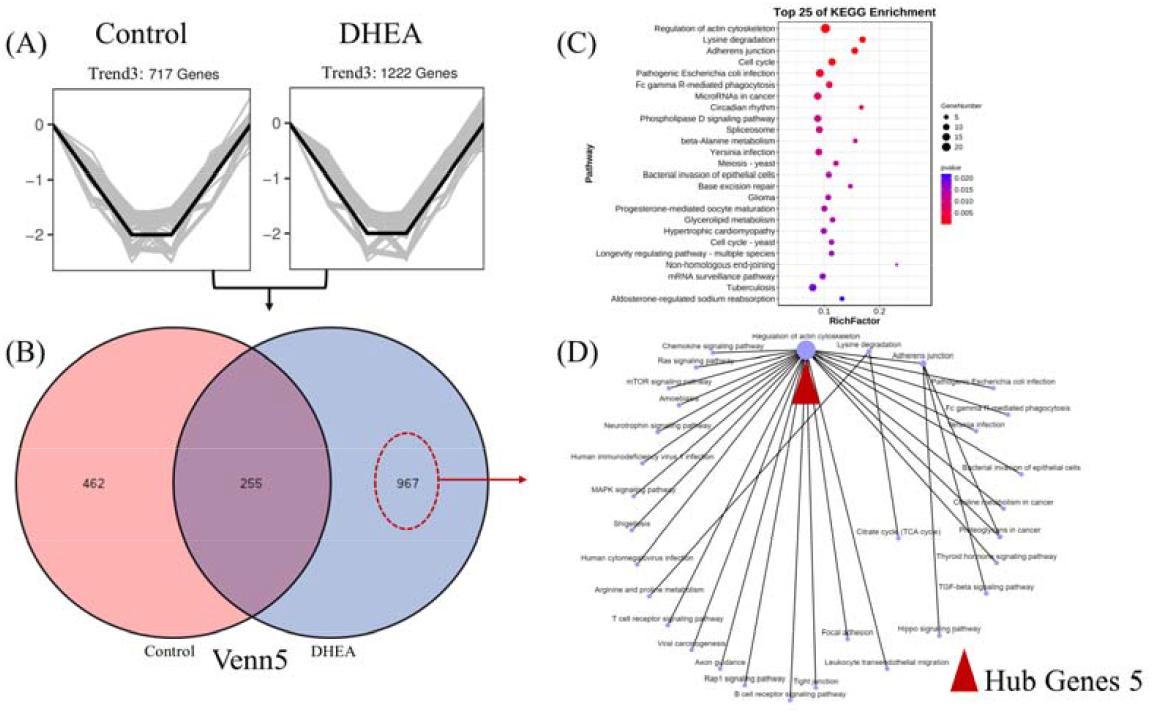
Analysis of genes expressed as ‘trend3’ in both groups. (a) 717 genes and 1222 genes were down-flat-up-regulated in each group. (b) 967 genes were abnormally down-flat-up-regulated in DHEA group. (c) KEGG analysis of the 967 genes abnormally down-flat-up-regulated in DHEA group. (d) Gene regulatory network diagram of the 967 genes abnormally down-flat-up-regulated in DHEA group.

**Figure10.**
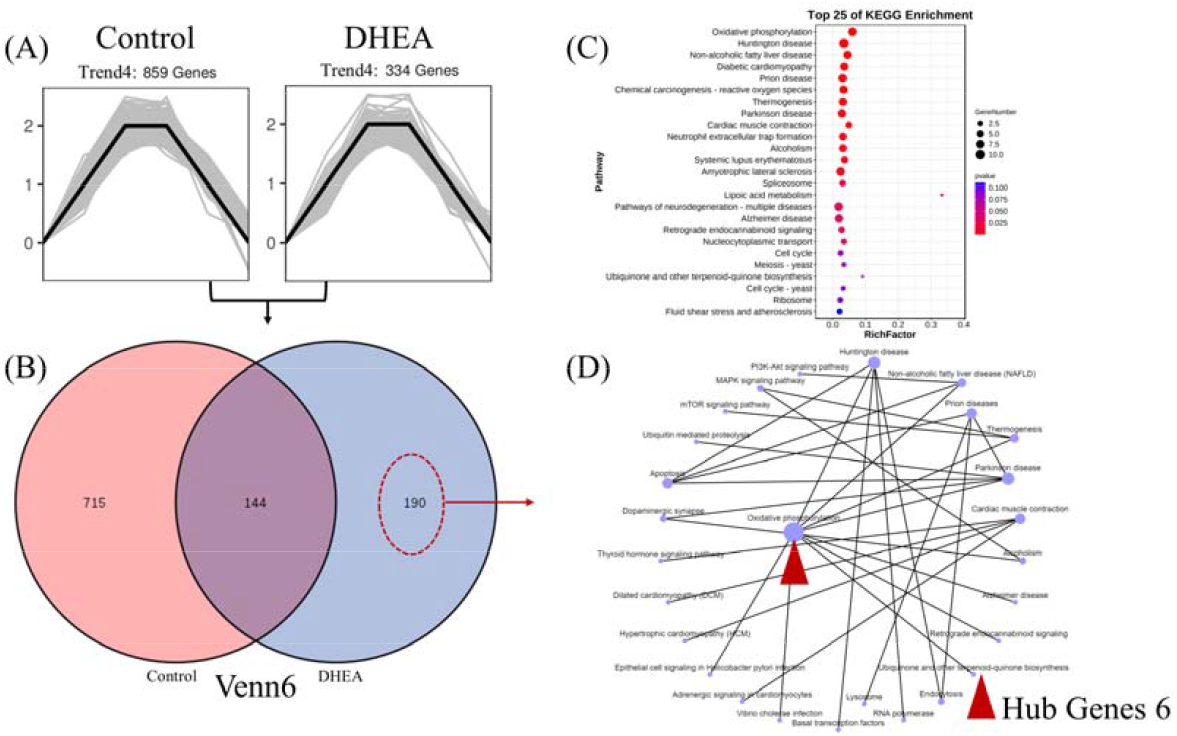
Analysis of genes expressed as ‘trend4’ in both groups. (a) 859 genes and 334 genes were up-flat-down-regulated in each group. (b) 190 genes were abnormally up-flat-down-regulated in DHEA group. (c) KEGG analysis of the 190 genes abnormally up-flat-down-regulated in DHEA group. (d) Gene regulatory network diagram of the 190 genes abnormally up-flat-down-regulated in DHEA group.

### 3.5 Genes clustering and KEGG analysis

KEGG analysis was used to detect the biological functions of the above six groups of selected genes. Genes abnormally up-regulated in the DHEA group were mainly focused on oxidative phosphorylation, thermogenesis, and the Parkinson disease pathway (Figure 5-10C). Genes abnormally down-regulated in the DHEA group were mainly focused on the cell cycle pathway. Genes abnormally up-regulated in the control group but down-regulated in the DHEA group were mainly focused on p53 signaling, PI3K-Akt, and glycolysis gluconeogenesis pathways. Genes down-regulated in the control group but up-regulated in the DHEA group were mainly focused on oxidative phosphorylation, central carbon metabolism in cancer, and the Alzheimer’s disease pathway. Genes abnormally down-flat-up-regulated in the DHEA group were mainly focused on the regulation of actin cytoskeleton pathway. Genes abnormally up-flat-down-regulated in the DHEA group were mainly focused on oxidative phosphorylation, apoptosis and Huntington disease pathways. The above results suggested that the most important pathways were oxidative phosphorylation and the cell cycle.

### 3.6 Gene regulatory network and hub genes

The selected genes from Venn diagrams were further investigated using gene regulatory network analysis to define the weight of different pathways. The result shows that pathways with the highest weight and strongest correlation were also oxidative phosphorylation and the cell cycle (Figure 5-10D). Some representative genes from a critical pathway were listed as potential hub genes (Table 5) and ranked by FPKM. The most significant difference was focused on mitochondrial oxidative phosphorylation. Hub genes abnormally up-regulated in the DHEA group included mtDNA coding subunits (mt-Nd1, 2, 3, 6 and mt-Co1,2), cytochrome oxidase (Cox8a, 7a2l, and 4i1), NADH-ubiquinone oxidoreductase (Ndufa1, 2, 6; Ndufb 9), and Uqcrq. Hub genes abnormally down-regulated in the DHEA group mainly focused on the cell cycle, which included Skp1, Ccnb1, origin recognition complex (ORC) subunit 1, and 5, Wee2, Mapk3, and Cdc20. All the other specific genes are listed in the supplementary documents (Table Venn1-6).

### 3.7 Immunofluorescence staining of Ptges3

According to the results above, Ptges3 was the top1 DEGs at time point1(Table1). The relative fluorescence intensity level of Ptges3 was significantly higher in DHEA group versus control oocytes at time point1(Figure11), which is consistent with the RNA-seq result.

## 4. Discussion

This study performed transcriptome sequencing throughout the course of PCOS development in a mouse model of adolescent PCOS to find the critical genes or pathways that regulate oocyte growth and quality. Identified DEGs in oocytes from the PCOS model were mainly focused on oxidative phosphorylation and the cell cycle.

Some mitochondria-related genes were abnormally and consistently up-regulated in the DHEA group, even after the DHEA injection regime was completed. Mt-Nd-x are key protein subunits of mitochondrial complex I, which is the main entrance for electrons to the respiratory chain, and its dysfunction will lead to decreased ATP production [17]. Mitochondrial complex I deficiency is associated with many clinical conditions, such as Leber hereditary optic neuropathy, mitochondrial encephalomyelopathy, and lactic acidosis [18]. Cytochrome c oxidase (COX) is the terminal enzyme of electron transport chains in mitochondria, and COX8a is required for maintenance of the structural stability of COX monomers and dimers [19]. COX7a2l is one of the three COX7a subunits, responsible for organization of the mitochondrial respiratory chain and preventing metabolic exhaustion [20]. Uqcrq is one of the ten nuclear genes encoding proteins of mitochondrial complex III, and mutations in Uqcrq are associated with a series of rare neuromuscular symptoms [21]. NDUFA-x encodes accessory subunits of mitochondrial complex I, and deficiency of this complex is the most common defect of the oxidative phosphorylation system. Mutations in NDUFA2 cause leukoencephalopathy [22], and patients with Leigh syndrome were found to harbor a homozygous mutation in NDUFA1[23]. Diseases caused by mitochondria deficiency may have some undetermined commonalities with PCOS, and the down regulation of mitochondrially encoded subunits of respiratory chain complexes may play an important role in the mitochondria dysfunction associated with PCOS [24].

Mitochondrial dysfunction has been proven to be a vital factor of many cardiovascular diseases and also plays an important role in the pathogenesis of PCOS [25]. Mitochondria convert nutrients into energy and release reactive oxygen species, but high levels of reactive oxygen species cause damage to mitochondrial components or even induce apoptosis. Growing evidence indicates that patients with PCOS have an increased level of oxidative stress and decreased levels of antioxidants [26]. The abnormal concentration of reactive oxygen species and antioxidant factors in follicular fluid from patients with PCOS may have adverse effects on oocyte quality, fertilization and embryo development [27]. mtDNA copy number is lower in women with PCOS and is negatively correlated with insulin resistance levels while positively associated with sex hormone-binding globulin levels [28]. However, some studies suggested that mtDNA number was increased in patients with PCOS and this increase was regarded as a feedback response compensating for mitochondrial dysfunction. These paradoxical results may reflect different stages of disease manifestation. mtDNA copy number increases during the early stage of PCOS as a compensatory effect, but decreases with disease development as the mitochondria fail to sustain normal function [29].

The markedly down-regulated genes in the DHEA group mainly focused on cell cycle. S-phase Kinase-Associated Protein1 (SKP1) is a core component of the Skp, Cullin, F-box containing complex, an E3 ubiquitin ligase that participates in a variety of cellular processes [30]. It was reported that reduced SKP1 expression induced chromosomal instability in both benign and malignant tumors [31]. Furthermore, SKP1 maintained synapsis in meiosis of both sexes [32], and also affected the balance of epigenetic regulation [33]. ORC proteins can bind DNA replication origins to initiate DNA synthesis, and ORC is especially important in controlling centriole and centrosome copy number [34]. Wee2 is an important oocyte-specific kinase, and its downregulation in oocytes resulted in high maturation-promoting factor activity and failure to mature to the meiosis-II stage [35]. Cyclin B1 is responsible for the composition of maturation-promoting factor, and Ccnb1-null oocytes completed meiosis I but arrested at the meiotic interphase [36]. Dominant or recessive CDC20 mutations have been linked to adverse oocyte maturation, and may act as a potential molecular markers of oocyte/embryo quality [37]. The relationship between all the hub genes mentioned above and PCOS has not been investigated until now. From these findings we propose that infertility or adverse pregnancy outcomes of patients with PCOS may result from mitochondrial dysfunction or abnormal meiotic regulation of oocytes.

At the first week of modeling, we found that Ptges3 was the top1 DEGs which was up-regulated in DHEA injection group at time point1. Ptges3 can act as an androgen receptor chaperone, which is present in many Hsp90 chaperone complexes and involved in assembling and disassembling some hormone or nuclear receptor/enhancer complexes[38]. Ptges3 and Hsp90 are of the basic minimal complexes required for the efficient folding and stabilization of steroid hormone receptors and essential for ligand responsive signaling[39]. In our research, Ptges3 was found up-regulated at the initial stage of modeling, which suggested it may play an important role in the early stage of PCOS.

## 5. Conclusion

This study provides a novel insight into differences in the transcriptomes of oocytes from control compared with PCOS mice. mtDNA-related genes and Cell cycle-related genes play the most important role in the development of PCOS. These findings could help focus on specific pathways or potential candidate genes that may play important roles during the development of adolescent onset PCOS. Ptges3 was the top1 DEGs which was up-regulated in DHEA group at the initial stage of modeling, which suggested it may play an important role in the early stage of PCOS.

## Supplementary Materials

The supplementary information can be downloaded at: https://figshare.com/.Table Venn1: Genes abnormally continuously up-regulated in the DHEA group (Figure 5B, marked by red dotted circle); Table Venn2: Genes abnormally down-regulated in the DHEA group (Figure 7B, marked by red dotted circle); Table Venn3: Genes abnormally down-regulated in the DHEA group (Figure 8B, marked by red dotted circle); Table Venn1 Venn4: Genes abnormally up-regulated in the DHEA group (Figure 9B, marked by red dotted circle); Table Venn5: Genes abnormally down-flat-up-regulated in the DHEA group (Figure 10B, marked by red dotted circle); Table Venn6: Genes were abnormally up-flat-down-regulated in the PCOS group (Figure 11B, marked by red dotted circle).

**Figure11.**
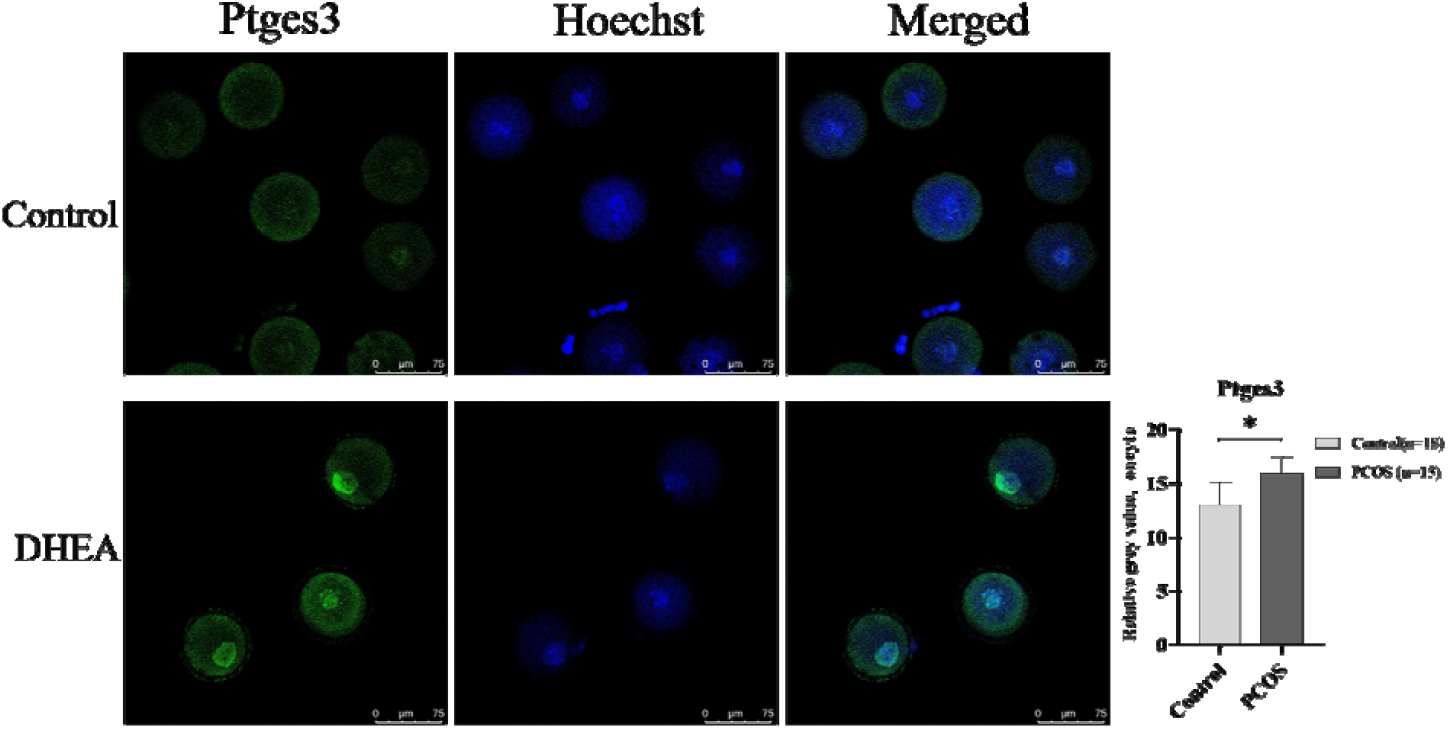
Immunofluorescence staining of Ptges3 in oocytes from control and DHEA group at time point1

## Author Contributions

1. Du Danfeng (First Author): Methodology, Software, Investigation, Formal Analysis, Writing-Original Draft.
2. Deng Ke (Co-First author): Data Curation, Investigation, Formal Analysis.
3. Fan Dengxuan: Resources, Supervision.
4. Xu Congjian (Corresponding Author): Conceptualization, Funding Acquisition, Resources, Supervision.

## Funding

This work was supported by funding from the National Natural Science Foundation of China to Xu Congjian (Grant number 82171639 and 82201807).

## Ethic Statement

All animal procedures were performed in accordance with the guidelines of Experimental Animals Management Regulations (version 2017-03-01, China).

Ethics approval was granted by the Institutional Animal Welfare and Ethics Committee Policies of Fudan University(approval number: 2020 Obstetrics and Gynecology Hospital JS-011 to Du Danfeng).

## Data Availability Statement

Raw data underlying this article are available in Figshare (https://figshare.com/). The authors affirm that all data necessary for confirming the conclusions of the article are present within the article, figures, and tables. Data analysis was performed using the OmicShare tools (https://www.omicshare.com/tools)

## Acknowledgments

We thank Liwen Bianji (Edanz) (www.liwenbianji.cn/) for editing the English text of a draft of this manuscript.

## Conflicts of Interest

The authors declare no conflict of interest.

## Notes

https://figshare.com/

### Competing Interest Statement

The authors have declared no competing interest.

### Summary of Updates

According to the data analysis result, immunofluorescence staining of Ptges3 was further performed and added to the manuscript(Figure11). All figures have been reordered in the manuscript.

